# Brain insulin signaling restores deficits in striatal dopamine release in overweight male mice with preexisting low D2-receptor expression

**DOI:** 10.1101/2025.10.28.685173

**Authors:** Miriam E. Bocarsly, Jacqueline B. Mehr, Evan S. Swanson, Sindhu Sriramoji-Virdi, Michael E. Authement, Sannidhi Shashikiran, Hannah C. Goldbach, Aya Matsui, Rachele Rimondini, Roland Bock, Veronica A. Alvarez

## Abstract

Obesity is characterized by insulin resistance, motivational impairments, and, in some cases, reduced availability of dopamine D2 receptors in the brain. However, whether the low D2 receptor levels represent a predisposing factor or a consequence of obesity, and how these processes are mechanistically linked, remains unclear. Here, we directly tested this causal relationship by selectively reducing D2 receptor density in striatal neurons. Male, but not female, mice with a low density of striatal D2 receptors consumed more food, gained more weight, and developed metabolic features of peripheral insulin resistance despite being maintained on standard chow. Motivational deficits preceded weight gain, manifesting as delayed circadian locomotor onset, reduced physical activity, and diminished effort to obtain food. In the brain, male mice with low D2 receptor density showed reduced dopamine release capacity and age-dependent alterations in brain insulin sensitivity. Prior to weight gain, brain insulin responses were blunted compared to those of controls, in which insulin potentiates dopamine release and enhances striatal acetylcholine signaling. Once overweight, however, these mice exhibited brain insulin hypersensitivity, with insulin strongly restoring dopamine release capacity. Together, these findings demonstrate that low striatal D2 receptor density predisposes male mice to an obesity-like phenotype through early dopaminergic dysfunction that precedes weight gain and is later compensated by insulin hypersensitivity in the brain.

## MAIN

Controlling body weight becomes increasingly difficult with advancing age. Even modest energy imbalances, arising from either excessive calorie intake, reduced physical activity, or both, accumulate to accelerate weight gain over time. Some individuals are particularly susceptible to these imbalances and subsequent development of obesity, yet the biological mechanisms that confer this vulnerability remain incompletely understood.

In humans, obesity has been associated with reduced striatal availability of dopamine D2 receptors^1,2^. Similar inverse associations have been reported in rodent models of diet-induced obesity^3–5^. Findings, however, are heterogeneous across striatal subregions and may be influenced by age^6–8^. A recent large-scale PET study addressed these discrepancies by examining adiposity-D2 associations with two tracers that differ in affinity and binding displacement by endogenous dopamine^9^. Using raclopride, a commonly used, relatively low-affinity D2-like receptor antagonist, the authors observed a negative association between adiposity and D2-receptor binding. However, the association was not evident when the same participants were scanned with fallypride, a higher-affinity tracer that is less sensitive to displacement by endogenous dopamine. This pattern suggests that changes in striatal dopamine tone, rather than receptor density per se, may underlie links between adiposity and D2 receptor signaling. Collectively, current evidence indicates a complex role for striatal dopamine and D2-like receptors in regulating body weight, feeding, and physical activity, underscoring the need for future mechanistic work.

Whether altered D2 receptor signaling is a cause or a consequence of weight gain remains a central question. Many PET studies are consistent with D2 receptor/dopamine changes arising secondary to obesity ^5,10^. Conversely, clinical observations in individuals treated with antipsychotic medications that are D2-receptor antagonists (e.g., clozapine, olanzapine) and are well known to promote weight gain, support a causal contribution^11,12^. Genetic evidence also aligns with this view: the TaqIA A1 allele of the DRD2 gene, associated with lower D2-receptor expression, correlates with higher BMI^13^. Together, these lines of evidence point to a bidirectional regulation between D2-receptor signaling and body weight.

Candidate mechanisms for this bidirectional link have been offered by preclinical studies. Experimenter-induced reductions of D2 receptor function decreased spontaneous and motivated physical activity, particularly voluntary wheel running, without necessarily altering body weight on standard chow or high-fat diet alone^3,14^. When mice were given simultaneous access to a high-fat diet and running wheels, control animals increased wheel running and limited weight gain, whereas mice with reduced D2 receptor expression engaged the wheel far less and became obese. These findings suggested that preexisting D2 receptor downregulation may increase vulnerability to obesity by constraining physical activity, thereby removing a behavioral buffer against excess energy intake.

An alternative explanation for the decreased physical activity may involve metabolic dysregulation arising from altered peripheral and central insulin signaling and its interactions with D2 receptors^15^. Peripheral D2 receptors in the pancreas regulate insulin secretion and glucose homeostasis, with downstream effects on body-weight regulation^16–18^. Beyond its peripheral actions, insulin also acts in the brain: insulin receptors are widely expressed, and within the striatum, insulin can potentiate dopaminergic signaling^19–21^. Dysregulation of insulin response, either peripheral, central, or both, could contribute to the mechanism by which low D2 receptors promote obesity risk with age

In this study, we built on the existing evidence and further identified the signaling mechanisms and circuitry that link D2 receptors to weight gain. We uncover an insulin-dependent mechanism connecting low D2 receptor function to obesity risk, more prominent in males than in females. In mice with reduced D2 receptor signaling, motivational impairments precede weight gain and are accompanied by diminished striatal dopamine release capacity and insulin brain response. Notably, once Drd2 haploinsufficient mice become overweight with age, the brain insulin response is potentiated and boosts dopamine release capacity to restore or overcome normal levels. These results integrate peripheral and central mechanisms to explain how D2 receptor hypofunction increases vulnerability to obesity and suggest that targeting insulin–dopamine interactions may offer sex-specific strategies for prevention or treatment.

## RESULTS

### Male mice with preexisting low striatal dopamine D2-receptor gain more weight on a nutritive diet

Mice with single-allele deletion of D2 receptor gene targeted to striatal projection neurons (iSPN-Drd2HET) and littermate control (Drd2^loxP/wt^) mice were group housed and maintained on *ad libitum* standard rodent chow. Body weights were measured weekly for 14 months, starting at 4 weeks old. Male iSPN-Drd2HET mice gained weight at a faster rate than male littermate controls (slope 1.1 ± 0.1 vs 0.8 ± 0.1 g/week for iSPN-Drd2HET vs control between 4 and 21 weeks old). With time, weight accumulated to create a significant weight difference between males of each genotype (control: 39.6 ± 2.3 g vs iSPN-Drd2HET: 50.1 ± 1.6 g; t(20)=3.65, p=0.002). The faster weight gain was sex-specific, with female iSPN-Drd2HET mice showing similar body weights to littermate Drd2^loxP/wt^ controls (Figure 1B; control 33.4 ± 1.3 g vs iSPN-Drd2HET 35.5 ± 1.5 g; t(19)=1.06, p=0.30). Thus, male mice appear more sensitive to Drd2 haploinsufficiency than females with regard to body weight.

**Figure 1.**
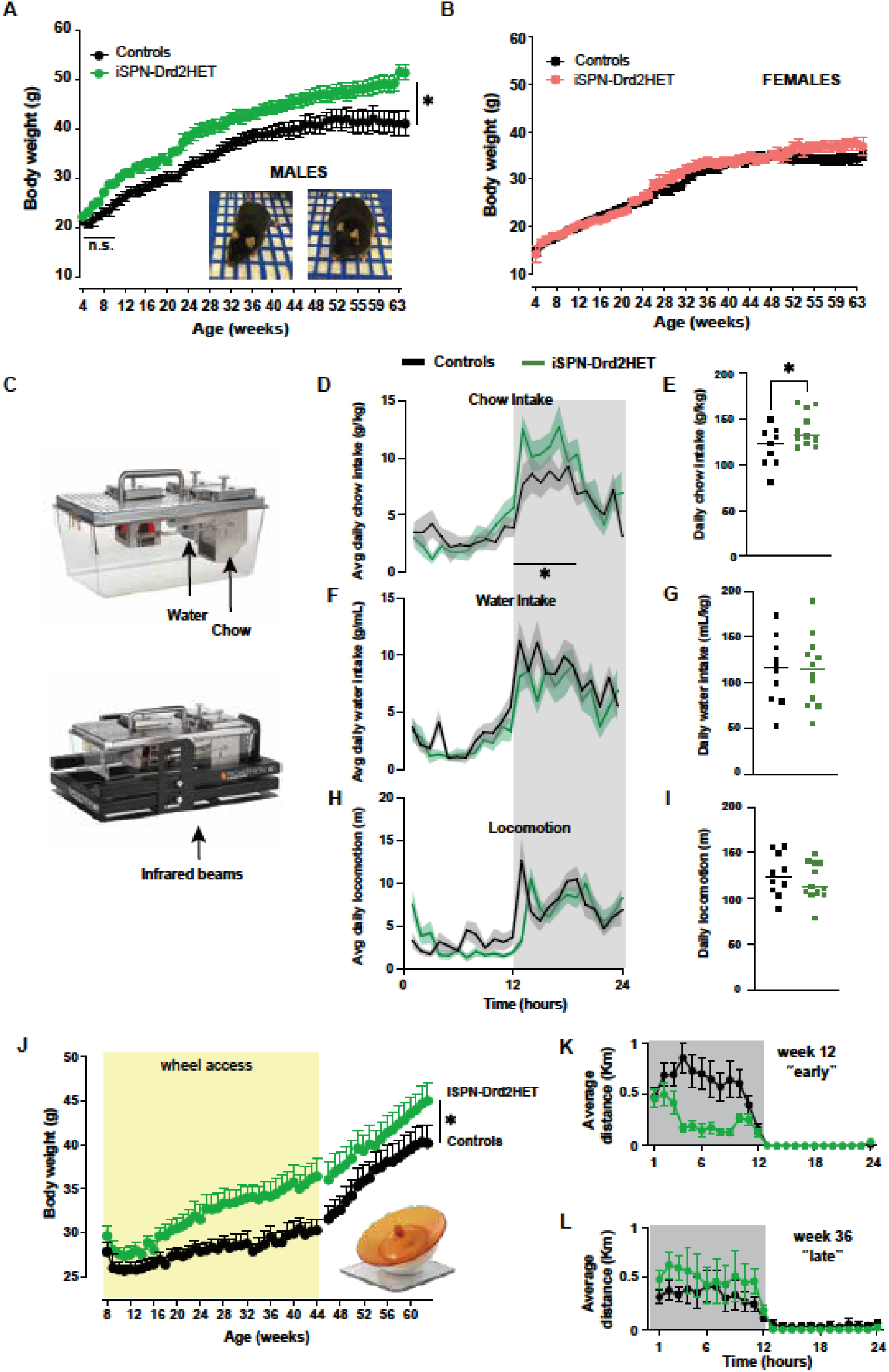
Male mice with reduced striatal D2 receptors exhibit age-dependent weight gain, increased food intake, and decreased physical activity. **A-B.** Longitudinal measure of body weight with age for male **(A)** and female **(B)** mice with knockdown of Drd2 gene in striatal projection neurons (iSPN-Drd2HET, green/pink) and littermate control (Drd2^LoxP/wt^, black). Symbols and lines are mean ± SEM. **C.** Top, Image of Sable System home cage (top) used to house mice for continuous high-resolution measurements of food and water intake. Bottom, Image with Sable System home cage with added infrared bean break detectors for assessing locomotor activity. **D, F, H**, Mean hourly measure of chow consumption (D), water consumption (F) and locomotor activity (H) for iSPN-Drd2HET (green) and control (black) male mice across the circadian light cycle. Lines representing mean and shaded are ± SEM. **E, G, I,** Mean daily values of chow intake (E), water intake (G) and locomotor activity (I) for iSPN-Drd2HET (green) and control (black) male mice. Symbols represent data from individual animals, lines represent group mean. **J.** Longitudinal measure of body weight for male mice housed with a running wheel from week 8 -44 (yellow) and after removal of running wheel for iSPN-Drd2HET (green) and control (black). Symbols and lines are mean ± SEM. **K,L**. Running distance per hour across the circadian light cycle for week 12 (K) and week 36 **(L)** for iSPN-Drd2HET (green) and control (black). Symbols and lines are mean ± SEM. For all panels, * denotes statistical significance p < 0.05.

Food, water, and locomotion were simultaneously monitored with high temporal resolution in Promethion Sable Systems for several weeks (Figure 1C). Male iSPN-Drd2HET mice (40 weeks old) showed circadian rhythms like controls but consumed more chow than controls (daily chow intake: 139 ± 5 g/kg for iSPN-Drd2HET vs 119 ± 7 g/kg for controls; t(18)=2.21, p=0.04; Figure 1D,E). No difference in water consumption was seen between genotypes (t(18)=0.004, p=0.99; Figure 1F,G). Further, no differences in basal body temperature or a fasting re-feeding task were seen in the iSPN-Drd2HET mice relative to controls, suggesting no gross changes in metabolism (t(28)=0.69, p=0.49; Supplemental Figure 5A,B,D). Because D2 receptors are also expressed in the pancreas, we verified that our genetic manipulation was restricted to the brain by dissecting pancreatic islets and measuring Drd2 mRNA expression. No differences in pancreatic Drd2 mRNA levels were detected between iSPN-Drd2HET mice and littermate controls (Supplemental Figure 5C).

While there was no overall difference in locomotor activity between groups (t(18)=0.73, p=0.47), the onset of locomotion in male mice with low D2-receptor is delayed by 60 mins during the active dark phase (Figure 1H,I, Supplementary Figure 1 shows cross-correlogram with 60 min shift). Given that in littermate control mice food consumption and locomotion ramp up with the onset of the dark phase without delays, the selective delay in locomotion seen in mice with low dopamine D2-receptor suggests that they eat first and then increase their locomotor activity. Female mice were not assessed in these measurements because they lacked a phenotypic difference in body weight.

### Low exercise levels contribute to weight gain in mice with low striatal D2-receptors

To further assess the contribution of locomotor activity to weight gain in mice with low D2 receptors, a separate cohort of male mice from both genotypes was provided with a running wheel in the home cage from age 8 to 44 weeks. Having access to a running wheel amplified weight differences between genotypes (Figure 1J). By 10 months old, the body weight difference between genotypes was 6.8 ± 2.2 g when mice had access to a running wheel compared to 4.5 ± 2.8 g without access (t(14)=2.80, p=0.01). This could be explained by differences in the rate of weight gain when exposed to a running wheel. Male control mice slowed weight gain by more than 75% when given access to a running wheel, from 0.8 g/week to 0.18 g/week, while mice with low D2-receptors slowed down by 50%, from 1.1 g/week to 0.53 g/week. With running wheel access, the rate of weight gain was twice as fast in male iSPN-Drd2HET compared to male controls (weight gain = 0.53 g/week vs 0.18 g/week, respectively; F(1,278)=12.85, p=0.0004).

Furthermore, when the running wheels were removed 8 months later, control mice rebounded and weight gain increased to 0.49 g/week (from 0.18 g/week with running wheels). However, weight gain rate in mice with low D2 receptors was not affected by removing running wheels (with wheel: 0.53, without wheel: 0.51 g/week, (F(1, 250)=1.29, p=0.26). These results indicate that access to a running wheel enhanced body weight differences between genotypes.

The running distance was measured throughout the day and found to follow a circadian pattern in both genotypes. We averaged the 24 hs running distance per week and found differences between the genotypes. Mice with low D2-receptors run less distance per day than controls. The difference in running is particularly evident early on when mice were 3-4 months old and control mice ran significantly more than iSPN-Drd2HET (Figure 1K). By 8 months of age, and 6 months of wheel access, control mice decrease their running and genotypic differences are eliminated (Figure 1L).

### Food motivation deficit mirroring obesity phenotype

Adult male iSPN-Drd2HET and littermate control mice were trained in an operant task for sucrose reward and tested on a behavioral economic task to assess high-effort and hedonic sucrose consumption. Briefly, mice were trained daily in an operant chamber to press a lever to earn a 20 mg sucrose pellet reward with an increasing fixed-ratio schedule (FR1, 3, 5, 10, 20, 32) while in a fasted state (Figure 2A). Mice of both genotypes acquired the task in approximately 20 hours of total training. No genotype differences were observed in overall training time (t(20)=1.63, p=0.12) nor the average frequency of head entry into the food port (t(20)=1.27, p=0.22; Supplemental Figure 3). Importantly, under the fasting conditions, male iSPN-Drd2HET mice showed similar body weight than age matched male controls (Figure 2C, no interaction between feed status and genotype F(1,20)=0.49, p=0.49) and no main effect of genotype F(1,20)=0.84, p=0.37, but main effect of feed status F(1,20)=58.86, p<0.0001), which eliminates a weight-based explanation of difference in the behavioral economic testing.

**Figure 2.**
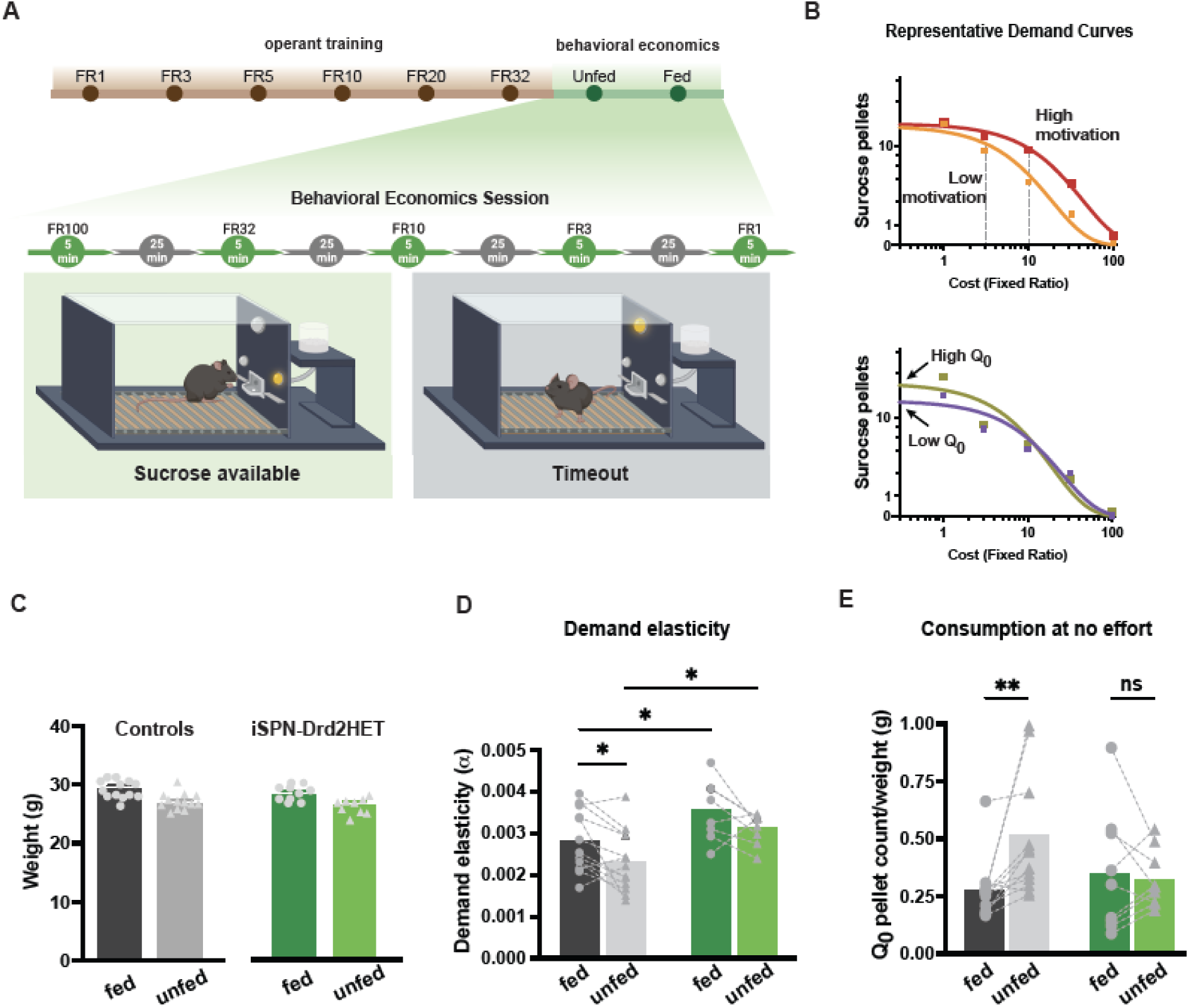
Impairments in motivational drive and interoception precede weight gain. **A.** Schematic depicting experimental timeline for operant training and setup within behavioral economics sessions **B.** Representative examples of demand curves for mice with different (*top*) motivation to earn sucrose pellets measured as changes in demand elasticity/alpha and (*bottom*) hedonic set points measured as Q_0_ when cost is null. Note that demand elasticity is inversely correlated with motivation. **C**. Bodyweight of young adult male mice at the start of behavioral training when mice were fed (solid) or unfed (shaded) for iSPN-Drd2HET (green) and littermate controls (black). **D, E**. Alpha and Q_0_ values extracted from demand curves for male iSPN-Drd2HET (green) and littermate control (black) mice tested in fed (solid) and unfed (fasted, shared) states. Bars represent mean; symbols represent data from individual animals. For all panels, *p < 0.05, **p < 0.01.

Once task-acquisition criteria were met, mice were run on the behavioral economics task in which intermittent access to sucrose pellets was given for 5 min in 30 min blocks with decreasing fixed-ratio schedules (FR100, 32, 10, 3, 1) in both fed and fasted states. From this task, demand curves were graphed and alpha and Q_0_, measures of motivation and consumption at null price, respectively, were extracted (Figure 2B).

As expected, consumption persisted to a higher fixed-ratio schedule (higher cost) when mice were fasted compared to fed, corresponding with a lower demand elasticity value (alpha) and indicative of greater motivation to seek food (Figure 2D). Furthermore, in control mice, the hedonic set point (Q_0_) was higher in a fasted state, consistent with the idea that hungry animals will consume more than satiated animals in effort-free feeding (Figure 2E).

Interestingly, male iSPN-Drd2HET mice showed motivational impairments towards food. Mice with low D2-receptor had higher demand elasticity values (alpha) than same sex controls, suggesting a lower motivation to seek sucrose (effect of genotype: F(1,20)=9.58, p=0.006). The higher alpha values of iSPN-Drd2HET mice were seen in both fed and fasted states (Figure 2; effect of metabolic state: F(1,19)=9.79, p=0.006). Furthermore, the hedonic set point (Q_0_) was not influenced by the hunger state in iSPN-Drd2HET mice. Both fed and fasted iSPN-Drd2HET mice showed similar Q_0_, indicating a similar amount of sucrose pellets consumed when hungry as when satiated (fed 0.35 ± 0.09, unfed 0.32 ± 0.04; t(8)=0.36, p=0.73). This may be from a subset of iSPN-Drd2HET mice that consumed a large number of pellets in the satiated state, possibly reflecting binge-like eating or insensitivity to metabolic cues.

### Low dopamine release capacity in the striatum of obese male mice with low D2R

Impairments in food motivation and exercise suggest alterations in dopamine transmission within the striatum. We used ex vivo fast-scan cyclic voltammetry to assess dopamine release capacity in striatal brain slices from male mice with low D2Rs and littermate controls (age = 17 ± 1.8 weeks; Figure 3A). Mice with low D2Rs showed decreased levels of evoked dopamine across a range of stimulation intensities compared to littermate controls (Figure 3B; interaction F(2.88,100.7)=3.53, p=0.02, main effect of stimulation F(2.88,100.7)=179.5, p<0.0001, main effect of genotype (1,35)=8.64, p<0.006). The mean dopamine transients have a peak amplitude of 1 ± 0.05 μM in control male mice versus 759 ± 29 nM for low D2-receptor male mice (t(74)=4.38, p<0.0001)). The decay of the dopamine signals was not significantly different, but there was a trend to slower times constant τ (0.26 ± 0.01 vs. 0.29 ± 0.02, t(29)=1.17, p=0.25 Figure 3B, inset). These findings indicate functional alteration in dopamine release capacity, and possibly dopamine clearance, which were detected in young adult mice (4 months) around the time of excess body weight gain.

**Figure 3.**
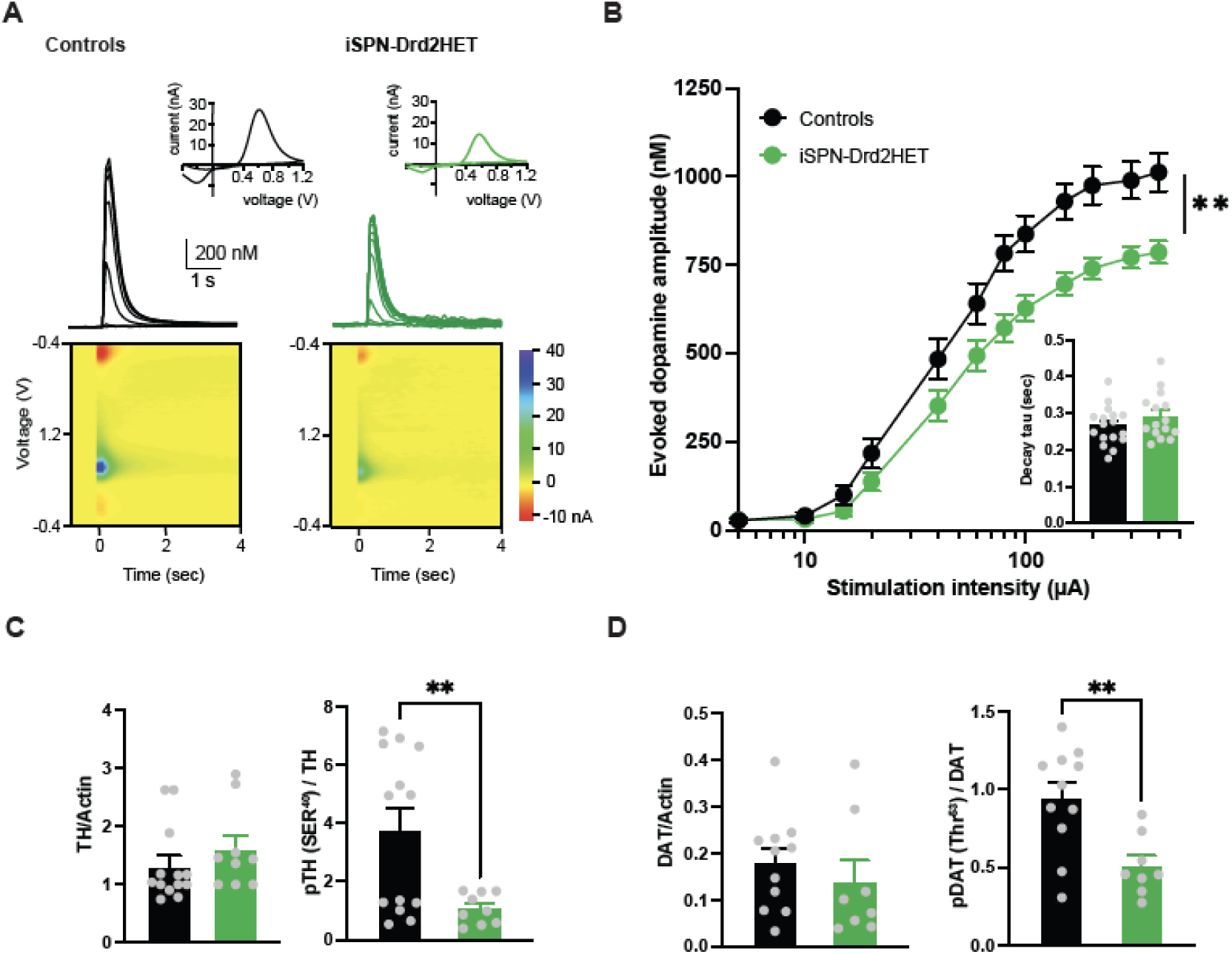
Dopamine release deficiency in mice with low striatal D2 receptors. **A.** Representative traces of evoked dopamine signals recorded in the striatum with FSCV in brain slices of control (black) and iSPN-Drd2HET (green) male mice. Superimposed transients were evoked by electrical pulses of increasing intensities. Inset, current-voltage plots of the peak signal show characteristic oxidation and reduction peaks for dopamine. Color plots show the time course (x-axis) of the current (color) vs voltage (y-axis) plots around the stimulation pulse delivered at t=0. **B**. Input-output curve shows mean amplitude of striatal dopamine signals as a function of stimulation intensity for control (black) and iSPN-Drd2HET (green) male mice (n = 33, 23 slices). Symbols and lines are mean ± SEM. **C**. Western blot quantification of protein levels for (*left*) total Tyrosin Hydroxylase normalized to total Actin and (*right*) phosphorylated serine40-Tyrosin Hydroxylase normalized to total TH in striatal tissue homogenate from control (black) and iSPN-Drd2HET (green) male mice. **D.** Western blot quantification of protein levels for (left) total dopamine transporter (DAT) normalized to total Actin and (*right*) phosphorylated Threonine53-DAT normalized to total DAT in striatal tissue homogenate from control (black) and iSPN-Drd2HET (green) male mice. Bars and lines are mean ± SEM; Symbols are individual sample data. For all panels, **p < 0.01.

We performed western blot analysis in tissue from the dorsomedial striatum to further examine molecular changes in key proteins involved in dopamine synthesis and reuptake. While overall levels of tyrosine hydroxylase were unchanged (t(20)=0.97, p=0.35) we found a decrease in the phosphorylation of this enzyme (t(20)=2.88, p=0.009), which is the rate-limiting step in the biosynthesis of dopamine (Figure 3C). The dopamine transporter (DAT), which is expressed in dopamine axons and clears dopamine from the extracellular space, also showed a selective reduction in the phosphorylated levels (t(18)=3.33, p=0.004) without impact on overall DAT levels between control and mice with low D2-receptors (Figure 3D; t(18)=0.92, p=0.37). Together, these functional and molecular analyses revealed reduced dopamine release capacity, likely driven by a combination of inhibited dopamine biosynthesis and reduced reuptake and recycling into vesicles. These alterations are likely contributing to the motivational impairments and reduced locomotor activity of male mice with low striatal D2R.

### Peripheral insensitivity to insulin in overweight male mice with low striatal D2-receptors

Obesity is characterized by profound metabolic changes, along with excess body weight and reduced motivation for food and exercise. We compared peripheral metabolic markers in male mice with low D2-receptor levels at two time points: early adulthood, corresponding to the onset of overweight, and late adulthood, after several months of sustained excess weight.

At 8 weeks of age, when mice are considered young adults, peripheral markers of metabolism were similar between male mice with low D2-receptors and littermate controls. Fasting blood glucose and insulin were no different between genotypes (Figure 4A,D) as well as no differences in their glucose tolerance when administered a bolus of glucose (1.5 g/kg i.p.; i.p.GTT) or insulin tolerance when administered a bolus of insulin (0.75 U/kg i.p.; i.p.ITT) compared to control littermates (Figure 4B,D). However, at 60 weeks of age, after several months of sustained weight gain, male mice with low D2-receptors showed increased fasting blood glucose levels (t(15)=2.21, p=0.04), impaired response to glucose on the i.p.GTT as reflected by a higher AUC (t(16.78)=3.93, p=0.001), decreased fasting blood insulin levels (t(16)=)2.27, p=0.03), and lower sensitivity to insulin on the i.p.ITT (t(5.56)=10.78, p=0002; Figure 4A-D). No differences were observed in fasting glucagon levels nor in a fast-refeeding task where iSPN-Drd2HET mice showed similar chow consumption and fluctuations in body weight during the fasting and refeeding than littermate controls (Supplemental Figure 5A,B,E). Female mice showed no differences in insulin or glucose measures at either time point or differences on the fasting and refeeding task (Supplemental Figure 4). Together, these data indicate that peripheral changes in insulin function in mice with low D2 receptors developed with time, and it suggests the peripheral insulin insensitivity is secondary to weight gain.

**Figure 4.**
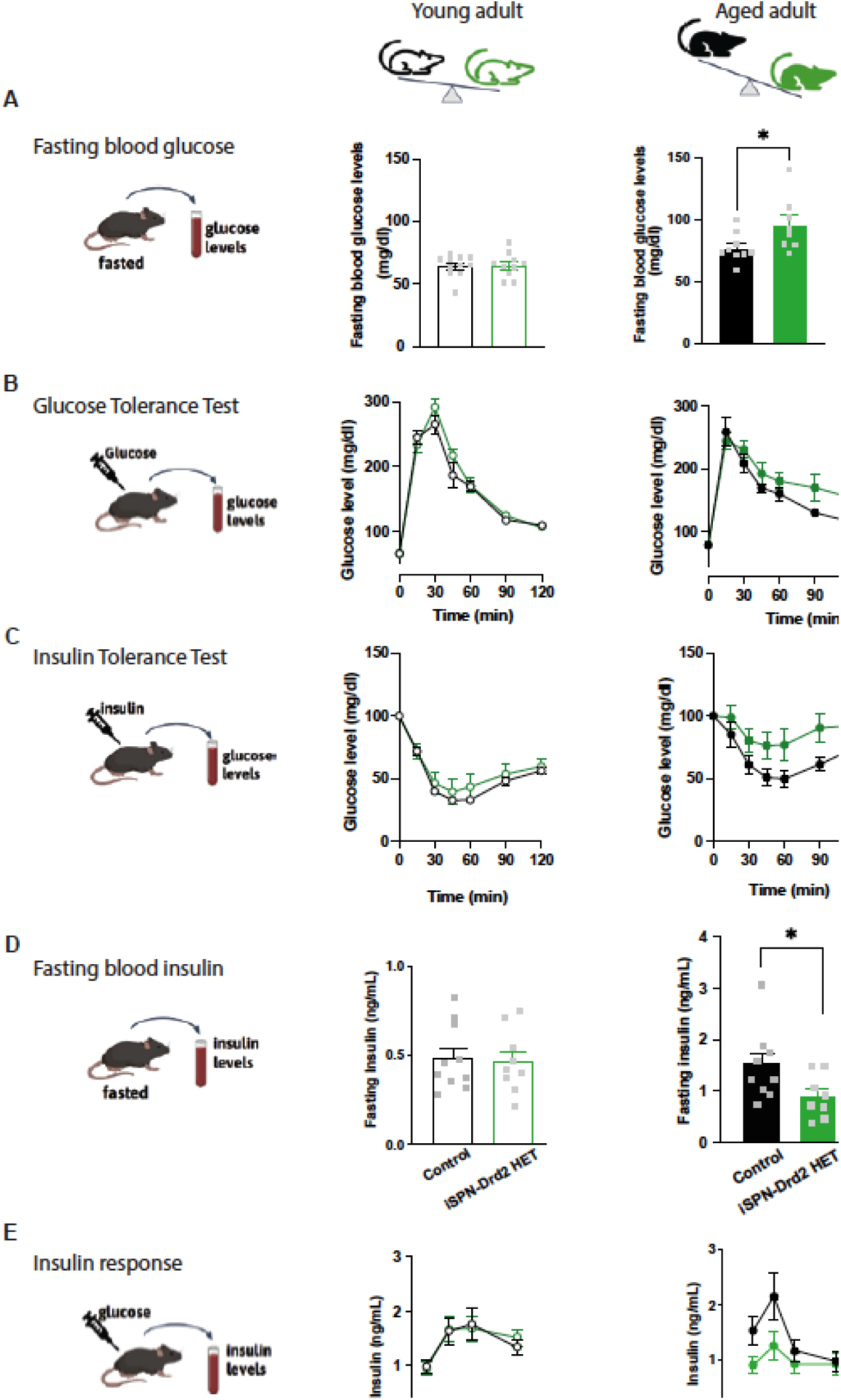
Metabolic signs of peripheral insulin resistance in aged overweight mice. All testing was performed in male young adult (8 weeks, left) and aged adult (60 weeks, right) iSPN-Drd2HET and control mice. **A.** Fasting blood glucose levels were determined. Symbols represent data from individual animals, while the bars show mean ± SEM. **B.** For the i.p. glucose tolerance test, mice were fasted overnight, administered an i.p. bolus of glucose and blood glucose was monitored for 2 hours. Again, male mice were tested as young adults (left) and aged adults (right). Lines represent mean ± SEM. **C.** Similarly, on the i.p. insulin tolerance test, mice were fasted, however, this time they were administered an i.p. bolus of insulin and blood glucose was monitored for 2 hours. As above, male mice were tested as young adults (left) and aged adults (right). Lines represent mean ± SEM. **D.** Fasting blood insulin levels were determined in both young (left) and aged (right) adult male mice. **E.** In a final test, fasted male mice were administered an i.p. bolus of glucose, and blood insulin levels were monitored for 1 h. For all panels, * denotes statistical significance p < 0.05.

### Insulin brain response involves acetylcholine and dopamine signals in the basal ganglia

To investigate how peripheral changes influence the brain’s response to insulin, we measured levels of two neuromodulators, acetylcholine and dopamine, in the basal ganglia of wildtype mice given subcutaneous insulin. We specifically examined the striatum, the principal input structure of the basal ganglia, which plays a key role in reward-motivated learning, tracking the value of cues and actions, and regulating feeding behavior.

Insulin application to mouse brain slices containing striatum was previously demonstrated to increase dopamine release via a mechanism that requires activation of cholinergic interneurons (references). Thus, we reasoned that systemic administration of insulin may increase acetylcholine in the striatum in vivo. Using expression of the acetylcholine sensor GRAB-ACh in the dorsal striatum and in vivo fiber photometry in head-fix fasted mice (∼ 10 months old), we found that insulin (0.3 U/kg, s.c.) increases the amplitude of spontaneous acetylcholine signals by more than 10% (t(8)=3.01, p=0.017, Figure 5A-B) compared to saline administration in the same animals. When compared to pre-injection amplitude, insulin but not saline administration significantly increased the amplitude of spontaneous acetylcholine signals in the striatum (one-sample test, saline: t(8)=1.71, p=0.125, insulin: t(8)=7.49, p<0.0001, Figure 5C). Thus, increasing peripheral levels of insulin leads to changes in acetylcholine signals in the striatum.

**Figure 5.**
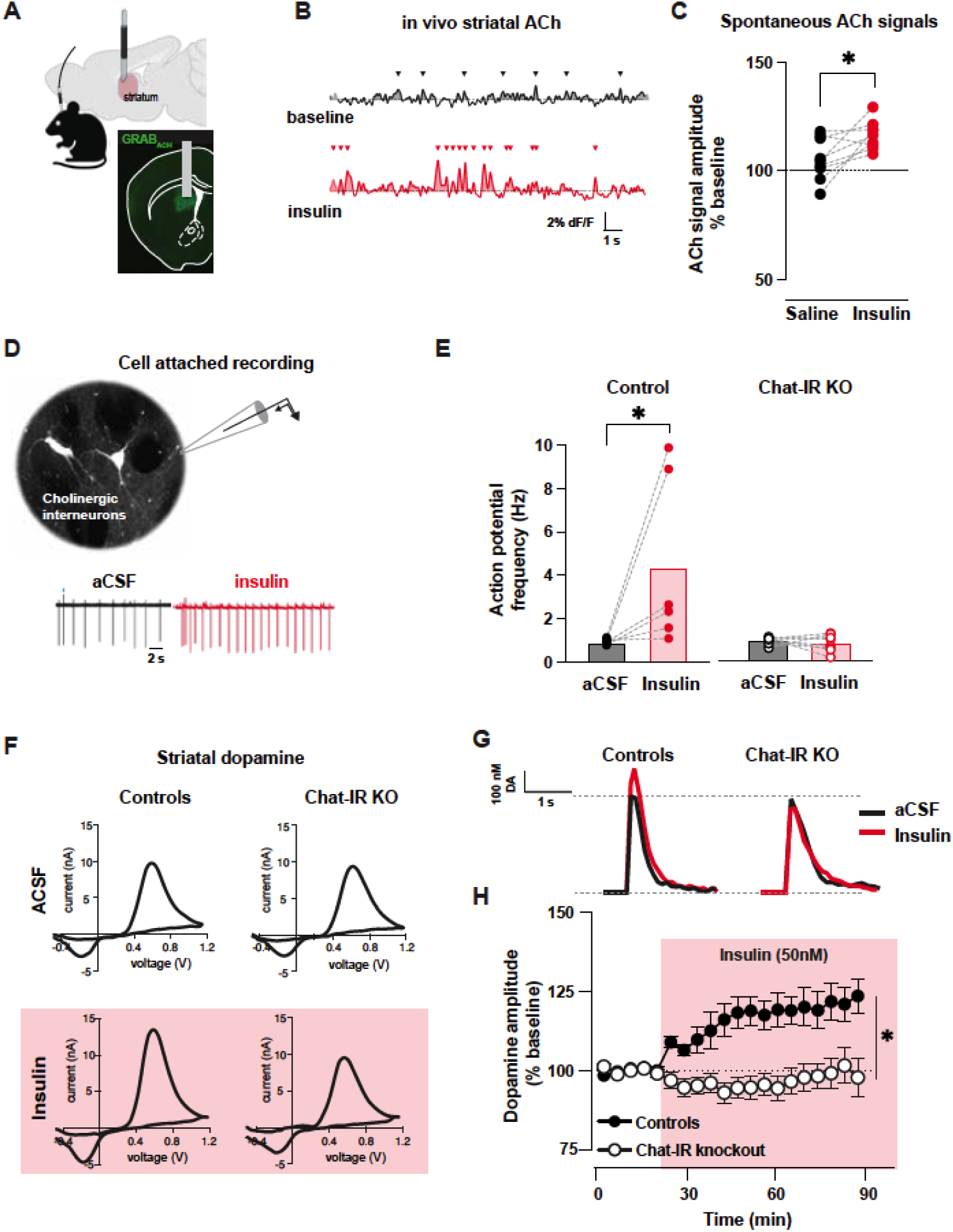
Insulin brain response involves amplification of striatal acetylcholine and dopamine. **(A)** In vivo fiber photometry of acetylcholine sensor in the dorsal striatum. Image of coronal brain section shows fiber optic placement (gray) and GRAB-ACh sensor (green) expression in the dorsal striatum. **B**. Representative examples of GRAB-ACh photometry signals showing spontaneous events (arrowheads) in mice before (black) and after insulin (red) administration (0.3 U/kg, s.c.). **C**. Amplitude of spontaneous acetylcholine signals after saline (black) and after insulin (red) normalized to the amplitude before administration. Symbols represent data from individual animals. **D**. Images of fluorescently labeled cholinergic interneurons in the dorsal striatum of ChAT-Cre reporter mice used for identifying these interneurons for electrophysiological recordings in brain slices. Representative traces of cell-attached recording from cholinergic interneurons showing the spontaneous firing of action potentials while in control solution (aCSF, black) and after bath application of 50 nM insulin (red). **E**. Frequency of action potential firing recorded in cholinergic interneurons before (black) and after insulin (red) in the striatum of control mice and mice with gene deletion of insulin receptors (IR) in cholinergic interneurons (Chat-IR KO). Symbols represent data from individual animals; bars represent group means. **F**. Striatal dopamine measurements in brain slices using fast-scan cyclic voltammetry produce representative voltammogram plots with the electrochemical signature of dopamine before and after bath application of insulin (50 nM, red shade). **G**. Time course of dopamine amplitude changes before and during insulin bath application in slices from control (solid) and ChAT-IR KO mice (open symbols). For all panels, data are presented as mean ± SEM, * denotes statistical significance p < 0.05.

This result is consistent with findings from the group of Rice et al. that showed insulin acts at receptors on striatal cholinergic interneurons to increase excitability and acetylcholine release which subsequently increases dopamine release.

We further proved this model *ex vivo* by assessing the effect of insulin administration on cholinergic interneuron firing, then perturbed the system via selective deletion of insulin receptors from cholinergic interneurons. Cholinergic interneurons are spontaneously active both *in vivo* and in the slice, allowing us to measure action potential frequency using cell-attached recordings in brain slices. In fasted control mice, insulin (50 nM) application increased the mean frequency of action potentials from 0.94 ± 0.20 Hz to 4.4 ± 1.5 Hz but had no effect in the firing of interneurons from fasted ChAT-IR knockout mice (Figure 5D,E; n=6-8 cells /genotype, interaction effect: F(1,12)=6.88, p=0.02, main effect of wash (F(1,12)=6.01, p=0.031, main effect of genotype: F(1,12)=1.06, p=0.025). We next measured downstream evoked dopamine signals using fast-scan cyclic voltammetry. In fasted control mice, insulin potentiated dopamine signals evoked by stimulation of the striatal tissue *ex vivo* (50 nM; Figure 5F-H). However, in fasted littermate mice with a deletion of insulin receptors in cholinergic interneurons (ChAT-IR knockout), the same concentration of insulin had no significant effect on evoked dopamine signals (interaction effect: F(2.55, 73.21)=6.15, p=0.002, main effect of insulin wash: F(2.55, 73.21)=6.45 p=0.001, and main effect of genotype: F(1,29)=13.5, p=0.001).

Thus, the brain response to insulin involves amplification of in vivo acetylcholine signals and enhanced excitability of cholinergic interneurons in the striatum. Insulin receptors in the striatal cholinergic interneurons are required for the insulin-mediated potentiation of striatal evoked dopamine.

### Overweight mice show insulin brain hypersensitivity that offsets dopamine deficiency caused by Drd2 haploinsufficiency

We next assessed the brain response to insulin in mice with low D2-receptors before and after the development of an overweight phenotype. Young (8-week) iSPN-Drd2Het mice show no difference in bodyweight relative to littermate controls, while old (60-week) iSPN-Drd2HETs are significantly heavier than controls when both are maintained on a standard chow diet (Figure 6A). In these mice, we measured insulin receptor expression levels using bulk mRNA quantification in tissue from the dorsomedial striatum. Insulin receptor mRNA levels were similar across genotypes in young adult mice (t(7)=0.18, p=0.86). However, in aged adult mice, higher levels of insulin receptor mRNA were found in the dorsomedial striatum of mice with low D2-receptors (Figure 6B; t(15)=2.14, p=0.04). This upregulation in insulin receptor expression led us to predict a stronger functional response to insulin in the striatum of iSPN-Drd2HET mice, which we tested by measuring insulin’s potentiation of striatal dopamine. Insulin (50 nM) increases evoked dopamine in slices from young and aged adult control male mice (insulin evoked dopamine increase in young: 24 ± 5% and aged: 19 ± 3%, t(13)=0.66, p=0.52; Figure 6C), consistent with findings from the experiment shown in Figure 5). In mice with low D2-receptors, the insulin response was age-dependent (t(36.24)=6.01, p<0.0001). As young adults, male iSPN-Drd2HET mice showed no potentiation by insulin (-5 ± 6% change in dopamine following insulin), while in late adulthood when mice were overweight, insulin produced a robust potentiation of dopamine that was even larger than in control mice (42 ± 5% change in dopamine following insulin; age x genotype interaction F(1,78)=19.01, p<0.0001, main effect of age: F(1,78)=13.26, p=0.0005, no effect of genotype F(1,78)=0.26, p=0.61, Figure 6C; time course in Supplemental Figure 6A-B). Importantly, in both genotypes the insulin potentiation was prevented by selective antagonists of the insulin receptor (HNMPA) and acetylcholine muscarinic receptor (scopolamine; genotype x drug interaction: F(2,100)=5.46, p=0.005, main effect of drug: F(2,100)=24.71, p<0.0001, insulin increased dopamine in both aged controls (p=0.03) and aged iSPN-Drd2HETS (p<0.0001), but not when co-applied with HNMPA or scopolamine (p values all >0.05, Figure 6D), indicating that similar mechanisms of insulin action are at play in control and Drd2 haploinsufficient mice.

**Figure 6.**
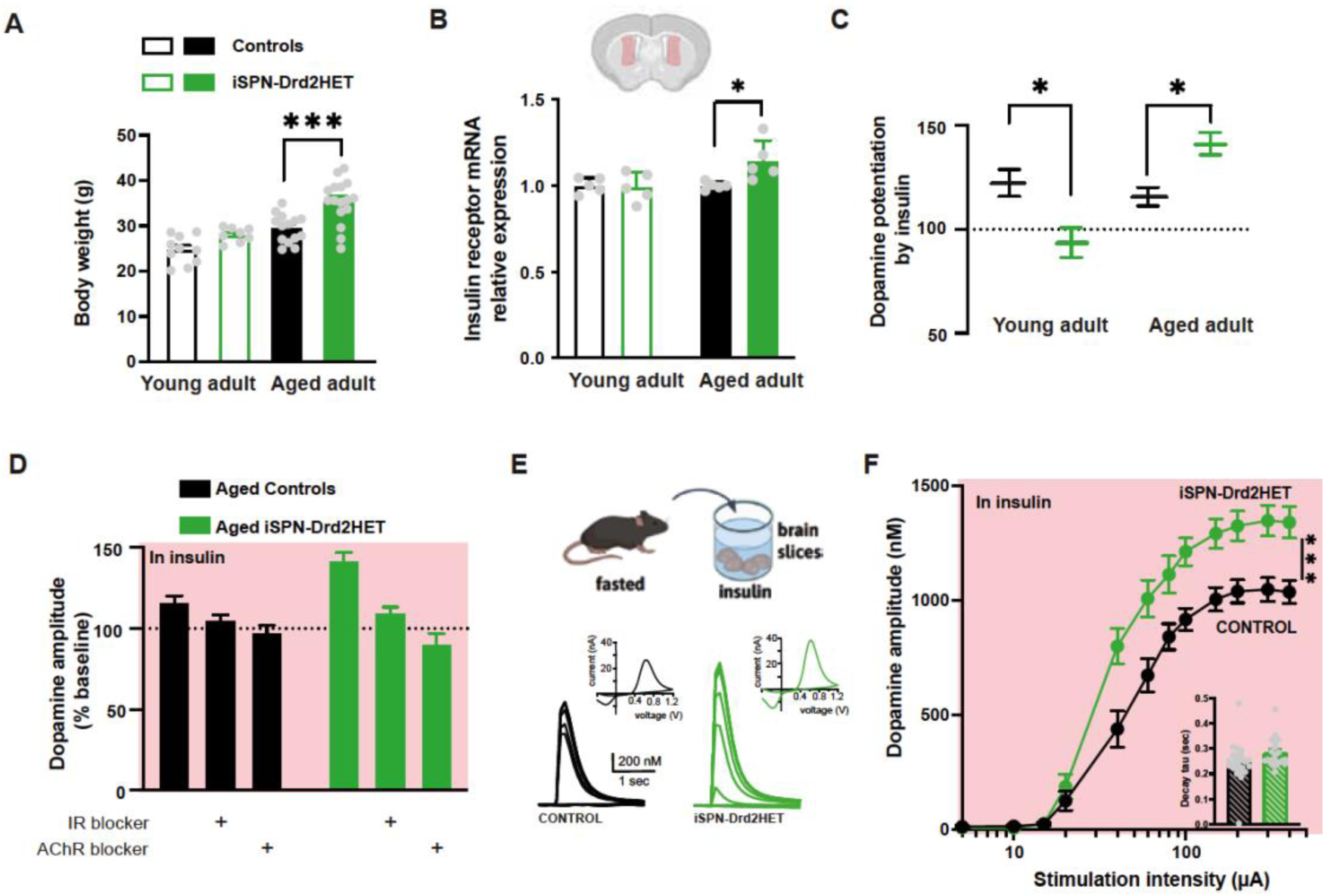
Hypersensitive brain insulin response develops in aged adults and overcomes dopamine deficiency seen in younger Drd2 haploinsufficient mice. **A.** Drd2 haploinsufficiency (iSPN-Drd2HET, green) leads to an overweight phenotype in male aged (64-week) adult (solid) mice compared to littermate control mice (black). Bars and lines are mean ± SEM; symbols are individual animal data. **B**. Insulin receptor mRNA expression in male striatal tissue from young adults (open bar; 8-12 weeks) and aged adults (solid bars, 64-68 weeks) control (black) and iSPN-Drd2HET (green) mice. **C**. Insulin-induced change in the magnitude of the evoked striatal dopamine responses measured by FSCV in brain slices from young adult and aged adult control (black) and iSPN-Drd2HET (green) mice. **D**. Insulin-induced change in striatal dopamine signals are blocked by an antagonist of the insulin receptor HNMPA and an antagonist of acetylcholine muscarinic receptor scopolamine (AChR blocker) in brain slices from aged adult control (black) and iSPN-Drd2HET (green) mice. **E**. Brain slices were prepared from overnight fasted aged adult mice and incubated in 50 nM insulin for 1 h (*top*) before measurements of evoked dopamine signals began with FSCV using stimulation pulses of increasing intensity (*bottom*) in control (black) and iSPN-Drd2HET (green). **F.** Input-output curve shows mean amplitude of dopamine signals evoked by electrical stimulation of increasing intensity in the dorsomedial striatum of control (black) and iSPN-Drd2HET (green) aged adult male mice. Symbols and lines are mean ± SEM. Inset, decay time constant of the dopamine signals relates to the dopamine reuptake properties in control (black) and iSPN-Drd2HET (green) mice. * denotes statistical significance p < 0.05, *** denotes statistical significance p < 0.001.

We hypothesized that this brain hypersensitivity to insulin might represent a mechanism for overcoming the deficit in dopamine release capacity seen in iSPN-Drd2HET mice. To test this hypothesis, we performed preincubation of the brain slices with insulin (50 nM) in fasted male mice from both genotypes and measured evoked dopamine signals in response to increasing stimulation intensity. After incubation with insulin, dopamine release capacity was larger in iSPN-Drd2HET mice compared to littermate controls at most stimulation intensities tested (interaction between genotype and stimulation intensity: F(2.88,128.5)=7.73, p=0.0001, main effect of stimulation intensity: F(2.88,128.5)=366.3, p<0.0001, main effect of genotype: F(1,45)=12.95, p=0.0008, Figure 6E,F). These results normalized the observations of young Drd2 haploinsufficient mice showing deficits in dopamine release capacity (Figure 3).

Altogether, we find that mice with low striatal D2 receptors have a blunted insulin brain response as young adults and display deficiencies in striatal dopamine and motivational drive. As aged adults, mice with this targeted Drd2 haploinsufficiency become overweight and develop insulin receptor upregulation and brain hypersensitivity to insulin. In overweight mice with low striatal D2 receptors, exogenous insulin is able to reverse and offset the dopamine deficiency, providing a possible mechanism for the brain’s dependence on insulin in these male mice.

## DISCUSSION

### Low Striatal D2 Receptor Density Predisposes to Metabolic Vulnerability

Mice with inherently low striatal dopamine D2 receptor density exhibited increased susceptibility to age-dependent weight gain and metabolic syndrome. Notably, this reduction in D2 receptor expression, induced from birth through targeted *Drd2* haploinsufficiency in striatal neurons, produced an obesity-like phenotype specifically in male mice. These findings establish a causal, sex-dependent relationship between striatal D2 receptor availability and vulnerability to obesity, independent of diet, as all animals were maintained on standard chow.

The mechanistic insights provided by this work highlight the insulin-dopamine interactions in the brain and its regulation of overeating and weight gain. A marked reduction in striatal dopamine release capacity preceded obesity onset and was accompanied by behavioral signatures of dopaminergic hypofunction, including diminished motivation, reduced physical activity, and impaired hedonic feeding. The emergence of an anhedonia-like phenotype before weight gain supports the idea that dopaminergic dysfunction is an initiating rather than a secondary event in the development of metabolic dysregulation.

### Behavioral and Motivational Consequences of D2 Receptor Deficiency

Reduced physical activity in *Drd2* haploinsufficient mice is consistent with prior studies showing that both global and striatal D2 receptor deletion from birth results in decreased exploration^3,14^. Extending these findings, we identified age- and sex-specific consequences of low striatal D2 receptor availability. Using high-precision behavioral monitoring, we found that male mice with reduced D2 receptor expression consumed more chow and exhibited delayed initiation of daily locomotor activity within the circadian cycle, indicating disrupted motivational and metabolic rhythms.

Our behavioral economics analyses further clarify this phenotype. Previous work using global D2 receptor knockdown or knockout mice reported an inverse correlation between D2 receptor levels and demand elasticity (α)^22,23^. Consistent with these observations, selective reduction of D2 receptors in striatal neurons increased effort-cost sensitivity (higher α values) and disrupted hedonic adjustment (Q₀), preventing behavioral adaptation between fasted and fed states. This combination of reduced motivation and impaired interoceptive regulation mirrors feeding behaviors commonly observed in obesity^24,25^.

### Brain Insulin Hypersensitivity as a Possible Compensatory Mechanism

In overweight male mice with low striatal D2 receptor density, we observed a state of brain insulin hypersensitivity characterized by elevated striatal insulin receptor expression and enhanced insulin-mediated potentiation of dopamine release. Exogenous insulin restored dopamine release deficits in these mice, suggesting a compensatory neuroendocrine adaptation. This brain insulin hypersensitivity may explain why sucrose pellet consumption was independent of the feeding state in our task. It suggests a persistent drive to elevate circulating insulin levels, possibly to boost dopamine release capacity, independently of metabolic state. This proposed mechanism aligns with previous evidence that striatal insulin signaling enhances striatal dopamine level and influences reward-driven feeding^26^.

Conversely, in young, lean male mice with reduced D2 receptor density, the brain’s insulin response was blunted before weight gain. This pattern aligns with the reward deficiency theory of obesity, which posits that individuals with diminished dopaminergic reward signaling overconsume food to compensate for reduced hedonic drive^27^. This brain insulin resistance that precedes weight gain and impairs dopaminergic circuits may represent a central vulnerability predisposing to overeating. Clinical evidence supports this model, as brain insulin resistance has been shown to alter dopamine turnover and contribute to depression and cognitive decline^28^.

Consistent with both clinical and preclinical evidence, females are generally less susceptible to peripheral insulin resistance^29^. Estrogen has been implicated as a key protective factor, and it may underlie the relative resilience to age-related weight gain observed in females in the present study. Elucidating the molecular and neuroendocrine mechanisms by which estrogen confers metabolic protection could provide valuable insight into sex-specific resilience and inform the development of targeted preventive and therapeutic strategies for metabolic disorders.

### Mechanistic Insights into Insulin–Dopamine Interactions

Our findings expand the mechanistic understanding of insulin’s actions in the basal ganglia. Previous work demonstrated that insulin potentiates dopamine release in both ventral and dorsal striatum through direct and indirect effects^19^. The indirect effect involves striatal cholinergic interneurons expressing insulin receptors^26^, whereas the direct effect acts through insulin receptors on dopamine axons regulating dopamine reuptake^19,30^.

We demonstrate that insulin receptors on striatal cholinergic interneurons are essential for insulin-induced dopamine potentiation. Cell-specific deletion of these receptors abolished insulin-driven increases in cholinergic firing and blunted insulin potentiation of dopamine release. Moreover, muscarinic receptor blockade prevented insulin’s effect, implicating muscarinic receptors in this process, in addition to nicotinic receptors^26^. Fiber photometry using a fluorescent acetylcholine sensor further revealed that insulin enhances spontaneous acetylcholine signaling in the dorsomedial striatum, confirming that insulin modulates both cholinergic and dopaminergic transmission.

In the midbrain, insulin also enhances reuptake but leads to suppression of somatodendritic dopamine release^31^. Although the suppression of somatodendritic release appears inconsistent with insulin’s potentiating of the axon terminal’s release within the striatum, these phenomena may be complementary. Dopamine release capacity at the axon terminals depends on the size and filling state of synaptic vesicle pools, which in turn rely on dopamine reuptake for vesicular refilling. Enhanced midbrain reuptake may therefore support sustained terminal release in the striatum under conditions of heightened insulin signaling.

### Bidirectional Interplay Between D2 Receptor Density and Overeating

The present findings provide strong evidence for a causal relationship between low density of striatal D2 receptors and the emergence of overeating, motivational deficits, and metabolic imbalance. Nevertheless, a bidirectional feedback mechanism remains likely. Overeating can reduce D2 receptor availability^24^, whereas caloric restriction increases it^5^. Human studies showing D2 receptor recovery following bariatric surgery^10^ underscore this reciprocity. Such a feedback loop could amplify dopaminergic and metabolic dysfunction: preexisting low D2 receptors heighten vulnerability to overeating-induced D2 downregulation, progressively weakening motivational drive and energy regulation.

Disentangling primary causal effects from secondary or compensatory adaptations will require temporally resolved and cell-type–specific interventions. We propose that brain insulin hypersensitivity represents a compensatory attempt to restore dopaminergic tone in the striatum. Within this framework, overeating may constitute a maladaptive effort to elevate circulating insulin and thereby normalize dopamine release in the D2 receptor-deficient brain.

### Integrative Perspective

Together, these findings delineate a mechanistic bridge linking dopamine signaling, insulin sensitivity, and metabolic regulation in male mice. Preexisting low density of striatal D2 receptor predisposes to motivational and metabolic dysregulation. Compensatory brain hypersensitivity to insulin initially mitigates dopaminergic insufficiency but may eventually contribute to maladaptive overeating. The resulting cycle of dopaminergic dysfunction, insulin dysregulation, and metabolic imbalance forms a self-reinforcing loop that accelerates obesity risk.

By establishing the causal sequence between dopaminergic deficiency, altered insulin signaling, and metabolic vulnerability, this study provides a framework for understanding the neurobiological basis of some forms of obesity. These insights highlight the intertwined roles of dopamine and insulin in energy balance and suggest new therapeutic avenues aimed at restoring dopamine–insulin homeostasis to prevent or reverse obesity in subjects with low D2 receptor density.

## METHODS

### Animals

All procedures were approved by the Animal Care and Use Committees of NIAAA, NIMH, and Rutgers University, and conducted according to specifications of the National Institutes of Health as outlined in the Guide for the Care and Use of Laboratory Animals. Male and female mice were group-housed on a 12 h light-dark cycle (lights on at 8:00 AM) and given *ad libitum* access to water and standard rodent chow, unless otherwise noted. Experimental procedures began at six weeks of age and weighed weekly.

All mouse lines used for breeding are commercially available. Mice with a targeted heterozygous deletion of the Drd2 gene resulting in Drd2 haploinsufficiency with validated knockdown of dopamine D2 receptor expression in striatal projection neurons are referred as iSPN-Drd2HET (Adora2a^Cre+/−^; Drd2^loxP/wt^)^32,33^. iSPN-Drd2HET mice were generated by crossing Drd2^loxP/loxP^ mice (B6.129S4(FVB)-*Drd2^tm^*^1^*^.1Mrub^*/J, JAX: 020631), which carry the conditional allele for Drd2, with Adora2a-Cre+/− mice (B6.FVB(Cg)-Tg(Adora2a-cre)KG139Gsat/Mmucd, MMRRC: 36158) which express Cre recombinase under the adenosine 2a receptor promoter. All mice were genotyped at weaning using real-time PCR with their respective probes by Transnetyx (Cordova, TN). To selectively remove insulin receptors from cholinergic interneurons, ChAT-ires-cre+/- mice (B6.129S-Chat^tm1^(cre)^Lowl^/MwarJ, JAX: 031661) were crossed with insulin receptor flox mice (B6.129S4(FVB)-Insr^tm1Khn^/J, JAX: 006955) generating Chat-IR KO mice. Cre-negative littermates in both groups were used as controls throughout the experiments. To identify cholinergic interneurons for recordings, ChAT-ires-CRE^+/-^ (B6.129S-Chatt^m1(cre)Lowl^/MwarJ, JAX: 031661) were crossed with Ai14 tdTomato mice (B6.Cg-Gt(ROSA)26Sor^tm14(CAG–tdTomato)Hze^/J, JAX:007914) to genetically label Cre+ neurons with red fluorescence.

### Metabolic Assays

#### Home-cage feeding, drinking, and locomotor activity

Mice were individually housed in the Mouse Promethion Core Behavioral System (Sable Systems International) home cages for two weeks with *ad libitum* access to both standard rodent chow and water. Each cage is equipped with a hanging food hopper filled with chow and water hopper connected to inverted laboratory balances, and an X, Y, and Z infrared beam break array. Mice were acclimated to housing conditions for at least 1 week prior to testing. Raw data were collected by SableScreen v3.3.12.0 (SableSystem) every second and extracted using Sable Systems Macro Interpreter v25.4.1 (SableSystem). Daily data presented is averaged daily, across mice during the second week of housing.

#### Glucose tolerance (ipGTT) and insulin tolerance tests (ipITT)

Mice were fasted overnight and injected with glucose (1.5 g/kg i.p.), followed by tail blood collection at 0, 15, 30, 45, 60, 90 and 120 minutes. Blood glucose levels were determined using the Elite glucometer (Bayer, Pittsburgh, PA). On a separate day, mice were fasted for 6 h before receiving insulin (0.75 U/kg, i.p.; Eli Lilly), and blood glucose levels were determined at the same intervals and using the same method as above. Tests were repeated in the same mice at 8 weeks (young adult) and 60 weeks (aged adult).

#### Insulin release testing

Mice were fasted for 6 hours and injected with glucose (1.5 g/kg i.p.), followed by tail blood collection at 0, 15, 30, and 60 minutes. Serum was collected and processed using the Ultra Sensitive Mouse Insulin ELISA Kit (CrystalChem, 62100) as per the manufacturer’s directions.

### Insulin Receptor mRNA Measurements

Quantitative polymerase chain reaction was used to measure relative mRNA levels in striatal brain tissue. Mice were anesthetized with isoflurane and decapitated. Brains were removed, and the striatum was dissected on ice using a 1 mm coronal matrix, placed in RNAlater, homogenized, and total RNA was purified using RNeasy Plus Mini kit (QIAGEN). cDNA was synthesized using iScript Reverse Transcription Supermix (Biorad). Actb (Mm01205647) and InsR (Mm00439688) TaqMan Gene Expression Assays (Applied Biosystems) were used to determine relative mRNA expression of the endogenous control gene β-actin, and the insulin receptor. Samples were run in triplicate and in parallel with negative controls using the StepOnePlus Real-Time PCR system (Applied Biosystems). The cycling conditions were: initial hold at 95°C (20 s), 40 cycles of 95°C (1 s) and 60°C (20 s). Relative insulin receptor mRNA levels were calculated using the ΔΔCt method.

### Western Blot

Mice were euthanized and dorsal striatum samples were collected and flash frozen. Tissue samples were lysed in RIPA buffer with protease inhibitor (Thermoscientific 89901), homogenized and then sonicated. Protein was evaluated using a Pierce BCA protein assay (Thermofisher 23225) at a 1:5 dilution. Protein (20 µg) was boiled in loading buffer (Bio-rad #1610374; 95°C, 7 min). Samples were loaded onto TGX stain-free mini gels (Bio-Rad #4561035) and run at 100 V for 2 h. Gels were transferred to a 0.2 µm nitrocellulose membrane using the Bio-Rad Trans-Blot Turbo at 2.5 amps, 25 V for 6 min. Poncu staining (Thermo Scientific A40000279) was utilized to confirm equal protein loading across wells. Membranes were blocked in 5% BSA in tris-buffered saline solution with tween (TBST) for one hour at room temperature with rapid shaking.

Primary antibodies (Anti-Tyrosine Hydroxylase:Millipore Sigma AB152; Anti-phospho-Tyrosine Hydroxylase: Millipore Sigma AB152; Anti-Dopamine Transporter: Proteintech 22524-1-AP; Anti-Dopamine Transporter: ABCAM ab183486; and Anti-Vinculin (Cell Signaling, 1390) were diluted at 1:1000 in TBST and incubated at 4°C with gentle shaking. Following primary incubation, membranes were rinsed with TBST 4×10 min with rapid shaking and then incubated with secondary HRP (Gt, Thermo Fisher, 18-4818-82 or Rb, Thermo Fisher, NA9340V, 1:5000) in 5% BSA in TBST for 1 h at room temperature with gentle shaking, followed by 2×15 min TBST rinses with rapid shaking. For development, membranes were covered in ECL (GE Healthcare, 17030484) for 2 min and imaged using chemiluminescence on the Chemi-doc imaging system. For blots in which multiple proteins could be detected from a single membrane, membranes were stripped between hybridizations with a stripping buffer (10% SDS, Tris HCl, pH 6.5, and 0.8% beta-mercaptoethanol) at room temperature for 15 min. Membranes were moved to TBST and rinsed 2×15 min. Membranes were re-imaged to ensure adequate stripping before re-blocking. Primary and secondary incubations were repeated as described. Band intensity was quantified using the Image Lab software (Bio-Rad).

### Voluntary Home-Cage Wheel Running

Mice were individually housed with *ad libitum* access to standard rodent chow and water. Mice were given unrestricted access to a running wheel (MedAssociates, ENV-047) for 36 weeks. Running distance was determined hourly, and body weights were taken weekly. Following 36 weeks of running, wheels were removed from the cage and body weights were tracked for an additional 16 weeks.

### Behavioral Economics Paradigm

Young adult (21±0.6 weeks old) male mice were run on an acquisition paradigm to earn a 20 mg sucrose pellet by pressing on a lever with an increasing fixed ratio schedule (FR1, FR3, FR5, FR10, FR20, and FR32). Each operant chamber was equipped with two levers, two cue lights, a pellet dispenser with food magazine, and a house light. Pressing the active lever resulted in the delivery of a single pellet according to the schedule. Presses to the inactive lever were recorded but had no scheduled consequences. The cue light above the active lever was turned on to indicate food availability. The house light was turned ON at the start of each session and turned back OFF at the end. The active lever side was counterbalanced across mice. Earned sucrose pellets were released into a port that had sensors to detect head entries. Criteria for moving to the next higher fixed ratio was as follows: earn 20 pellets for three consecutive training days and have at least twice as many active lever presses as inactive lever presses on the final of the three consecutive days. Mice were fasted overnight before each training session (food off between 4 and 6pm and sessions began between 9 and 11am).

Following completion of FR32 training, mice immediately moved to the behavioral economics (BE) paradigm. In BE, mice were similarly fasted overnight before each session. This paradigm consisted of five 5-min “food availability” bins with diminishing fixed ratio schedules (FR100, FR32, FR10, FR3, and FR1), indicated by an illuminated light only over the active lever, interspersed with four 25-min “food unavailability” bins, indicated by illumination of only the operant box house light. Consumption at each fixed ratio was graphed and demand curves were fitted according to Gilroy et al., inverse sine hyperbolic transformation^34^. From this fit, alpha and Q_0_ values were generated for each BE session.

Mice were run on BE for at least six sessions and until the 3 most recent sessions produced stable α and values (<25% variability). Following stabilization, mice were provided ad lib access to chow and tested on BE for three more days.

For data analyses, active lever responses in each food availability bin from the three final fasted BE sessions and three unfasted BE sessions for each mouse were averaged, and a demand curve was fit from these data points according to Gilroy et al^34^. Thus, average alpha and Q_0_ values were generated for each animal in an unfed and fed state.

### Fast-Scan Cyclic Voltammetric and Electrophysiological Recordings

#### Brain slice preparation

Male mice (17±1.8 weeks old) were fasted overnight and on the day of experiment, mice were deeply anesthetized with isoflurane, decapitated, and brains were rapidly harvested and placed in ice-cold cutting solution (in mM): 225 sucrose, 13.9 NaCl, 26.2 NaHCO_3_, 1 NaH_2_PO_4_, 1.25 glucose, 2.5 KCl, 0.1 CaCl_2_, 4.9 MgCl_2_, and 3 kynurenic acid. Brains were sliced coronally using a vibratome (240 μm) in cold cutting solution. Brain sections containing dorsomedial striatum were allowed to recover in oxygenated ACSF containing (in mM): 124 NaCl, 2.5 KCl, 2.5 CaCl_2_, 1.3 MgCl_2_, 26.2 NaHCO_3_, 1 NaH_2_PO_4_, and 20 glucose (∼310-315 mOsm) at 33⁰C for 30-60 min and maintained at room temperature after that until recording.

#### Fast-scan cyclic voltammetry recordings

Fast-scanning cyclic voltammetry was performed in the ventral aspect of the dorsomedial striatum from iSPN-Drd2HET and littermate control (Drd2^loxP/wt^) mice to determine levels of evoked dopamine. Carbon fibers (7μm diameter) were inserted into glass pipettes to create carbon fiber electrodes (CFEs), which were hand-cut to expose ∼150μm of the fiber past the pipette tip. Before use, CFEs were backfilled with 3M KCl internal solution. CFEs were conditioned before recordings with a triangular voltage ramp (-0.4 to 1.2 and back to -0.4V versus Ag/AgCl reference at 400V/s) delivered every 15ms. During recordings, CFEs were held at -0.4V versus Ag/AgCl while the same triangular voltage ramp was delivered every 100ms. Dopamine transients were evoked by a single pulse (0.2ms) of electrical stimulation administered every 2 minutes by a glass micropipette filled with ACSF that was placed near the CFE tip. To generate input-output curves, the intensity of electrical stimulations was varied (5, 10, 15, 20, 40, 60, 80, 100, 150, 200, 300, or 400µA) with each episode. Experiments were conducted in ACSF and maintained at 31-33℃. In a subset of experiments, slices were pretreated in an ACSF solution containing vehicle (BSA at 0.05-0.1mg/mL) or insulin (50nM, in BSA solution) for 60-90 minutes prior to recording; these slices remained in this solution for the duration of the experiment. Data were acquired and analyzed using a Chem-Clamp amplifier (Dagan) and I/O board (National Instruments) with a custom-written software VIGOR using Igor Pro (Wavemetrics) running mafPC (courtesy of M.A. Xu-Friedman).

#### Cell-attached electrophysiology recordings

Coronal brain slices (240 µm) containing the dorsomedial striatum were prepared from mice (males, 8-16 week-old, fasted overnight) with genetically-identified cholinergic interneurons (ChAT-ires-CRE^+/-^; Ai14 tdTomato). Cell-attached recordings in voltage clamp mode were performed to measure the spontaneous firing frequency of cholinergic interneurons. Electrodes were filled with filtered ACSF. A gigaohm seal was achieved, maintained and monitored. Recording from cells in which the gigaohm seal had degraded were excluded. Data were acquired at 5 kHz and filtered at 1 kHz using Multiclamp 700B (Molecular Devices). Data was analyzed using pClamp (Clampfit, v. 10.3).

### In vivo Fiber Photometry

#### Surgery

Acetylcholine sensor GRAB_ACh (AAV9-hSyn-Ach3.0(Ach4.3); titer: adjusted to 1×10^13^ vg/mL, WZ Biosciences) was injected unilaterally in the dorsomedial striatum (coordinates mm from bregma: +0.9 AP, ± 1.3 ML, −2.8 DV) of C57BL/6J mice. Immediately following viral vector injections, a custom headpost and optic-fiber cannula (400 μm; 0.37 NA; 2.5 mm long; Doric Lenses) were implanted above the injection site and secured to the skull with Metabond (Parkell). Mice were single housed after fiber implant surgery to prevent damage to the implants.

#### Photometry system

Thorlabs LED drivers (LEDD1B) were used to drive Thorlabs 405 nm (M405F1) and 470 nm (M470F4) fiber-coupled LEDs connected to a six-channel Doric filter cube (FMC6_IE(400-410)_E1(460-490)_F1(500-540)_E2(555-570)_F2(580-680)_S) by 0.5 NA / 400 μm Ø patch cords (Thorlabs, M301L01). Subject cords were custom-ordered from Doric (MFP_400/440/1100-0.37_1m_FCM-ZF1.25(FP)_LAF). Newport Femtowatt Receivers (Doric, NPM_2151_FOA_FC) were connected to the filter cubes using custom 0.5 NA / 600 μm Ø patch cords (Thorlabs). Femtowatt receivers fed into a Tucker Davis Technologies RZ5P processor. All fiber optic cords were photobleached for 24 hours before the start of the experiment. Data were collected and demodulated using Tucker Davis Technologies Synapse software.

#### Data collection and analysis

Mice (10-12 months old) were habituated to be head-fixed in an acrylic tube within a noise-attenuating chamber with dimmed light for the recording duration (30-60 min). Each experimental day, LEDs were calibrated daily to 20 μW (405 nm) and 30 μW (470 nm), as measured at the fiber tip. Data was collected at 1017 Hz and low-pass filtered at 4 Hz. On the experimental day, baseline photometry signals were recorded for 5 mins before mice were injected subcutaneously with saline and insulin in consecutive sessions. After injection, mice were kept in homecage for 30 minutes and headfixed after which photometry recordings were resumed for another 5 mins. Data were converted to ΔF/F signals using GuPPy^35^, an open-source Python package (RRID: SCR_02235345). The same Python package was used to detect spontaneous events and calculate amplitude. The preprocessed data were then downsampled to 100 Hz for further analysis and plotting in Python. Z-scored amplitudes of spontaneous events were calculated using the means and standard deviations of the entire combined baseline and treatment ΔF/F traces.

#### Drugs

Insulin (human source, Sigma-Aldrich I9278) was used in *in vivo* experiments (i.p.ITT and fiber photometry; s.c.); for *ex vivo* experiments, insulin (from porcine pancreas, Millipore-Sigma catalog I5523) was dissolved at 50 nM in ACSF with bovine serum albumin (0.05-0.1 mg/ml). Cell-permeable insulin receptor antagonist HMNPA-AM3 (trisacetoxymethyl ester; Enzo Life Sciences BMLEI2480005); acetylcholine muscarinic receptor antagonist scopolamine hydrobromide (Tocris 1414) was applied at 1 µM dissolved in ACSF. For i.p.GTT, Glucose (Sigma-Aldrich G8270) was administered (i.p.). All subcutaneous and intraperitoneal administration of drugs was delivered at 10 ml/kg body weight.

### Statistical analysis

Analyses were performed in Prism 10 (GraphPad), using two-sided tests. For behavioral economic data, outliers were identified using the Grubbs’ test. Data making three or more comparisons were analyzed using two-way ANOVA, with the addition of repeated-measures (RM) or a mixed-effects model, as appropriate. For RM-ANOVA, sphericity was assessed with Mauchly’s test, and Greenhouse–Geisser correction was applied when appropriate. Significant main effects or interactions were followed-up with pairwise tests corrected for multiple comparisons (Sidak or Tukey). Data comparing two groups (ex. average daily locomotion or intake, protein expression, metabolic parameters, etc. between genotypes) were compared with an unpaired Student’s t-test. Paired or independent sample t tests were used to analyze the remainder of data. Results were considered significant at an alpha < 0.05. All data are presented as mean ± SEM and individual animal data also showed whenever possible. Sample sizes were chosen based on previous research employing similar approaches and are sufficient for detecting strong effect sizes while using the minimal number of animals.

## ACKNOWLEDGEMENT

This research was supported by the Intramural Research Program of the National Institutes of Health (NIH) to VAA (ZIA MH002987 and ZIA AA000421) and to MEB (NIH AA027750). The contributions of the NIH authors were made as part of their official duties as NIH federal employees, are in compliance with agency policy requirements, and are considered Works of the United States Government. However, the findings and conclusions presented in this paper are those of the authors and do not necessarily reflect the views of the NIH or the U.S. Department of Health and Human Services.

## AUTHOR CONTRIBUTIONS

Conceptualization and Design: VAA, MEB; Methodology and Data Collection: MEB, JBM, JHS, HCG, ESS, RR, SSV, MA, AM; Data Analysis: MEB, JBM, HCG, MA, ESS, RB; Interpretation of Results: VAA, MEB, JBM; Manuscript preparation: VAA, MEB, JBM; Manuscript Review & Editing: all authors; Funding Acquisition: VAA, MEB.

## MATERIAL AVAILABILITY

This study did not generate new unique reagents.

## Data and code availability

The datasets supporting the current study have been deposited at Mendeley.com (DOI:10.17632/hgd3mkypwj.1) and are publicly available as of the date of publication. Original code was generated for acquisition of voltammetry data (VIGOR) made available from https://bitbucket.org/r-bock/vigor/; and code for behavioral economic task, available at public repository (https://bitbucket.org/r-bock/medpc-sa/). Any additional information required to reanalyze the data reported in this paper is available from the lead contact upon request.

