## Extended Data Figures for "Brain insulin signaling restores deficits in striatal dopamine release in overweight male mice with preexisting low D2-receptor expression"

Extended Data Figure 1. Bocarsly et al.

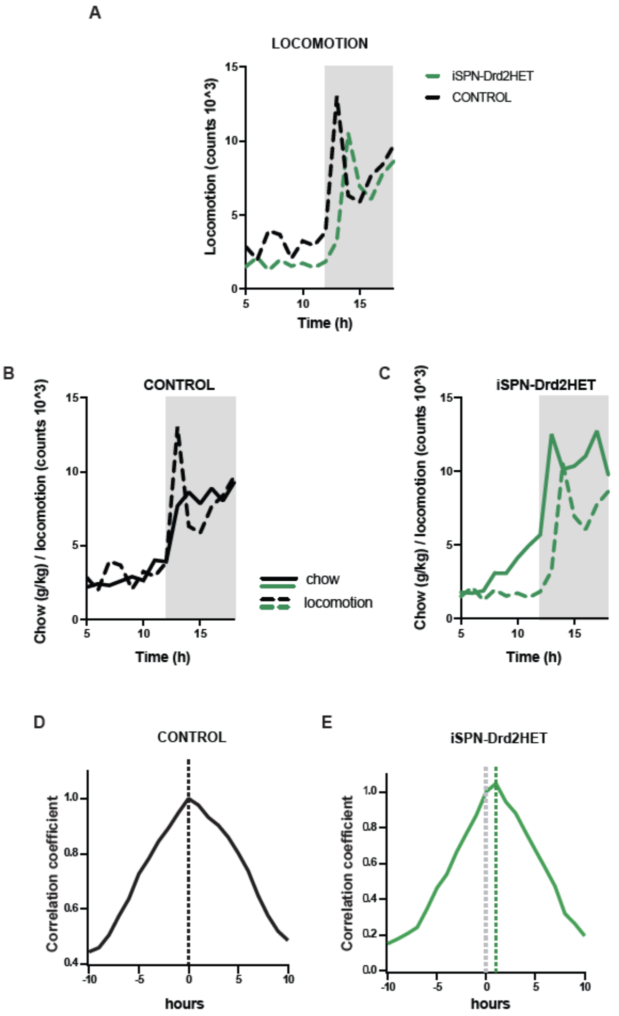


**Extended Data Figure 1. Delayed onset of circadian locomotor activation in male mice with low striatal D2 receptors**

**A.** Expanded timeline in the circadian cycle for the 24-h locomotor activity, centered around the light change from inactive (light) to active (dark, shaded) phase for iSPN-Drd2HET (green) and littermate control (Drd2$^{LoxP/wt}$, black) mice. Dashed lines are mean. **B-C**. Expanded timeline in the circadian cycle for the 24-h locomotor activity (dashed line) and chow intake (solid line) for **B**, littermate Drd2$^{LoxP/wt}$ control mice (black) and **C**, iSPN-Drd2HET mice (green). Lines are mean. **D-E** Cross-correlograms between locomotor activity and chow intake across the circadian cycle show **D**, the peak at 0 hours for control mice (black) and E, the peak at 1 hour for iSPN-Drd2HET mice (green), indicative of a time shift between the onset of feeding and locomotion.

**Extended Data Figure 2. Bocarsly et al.**

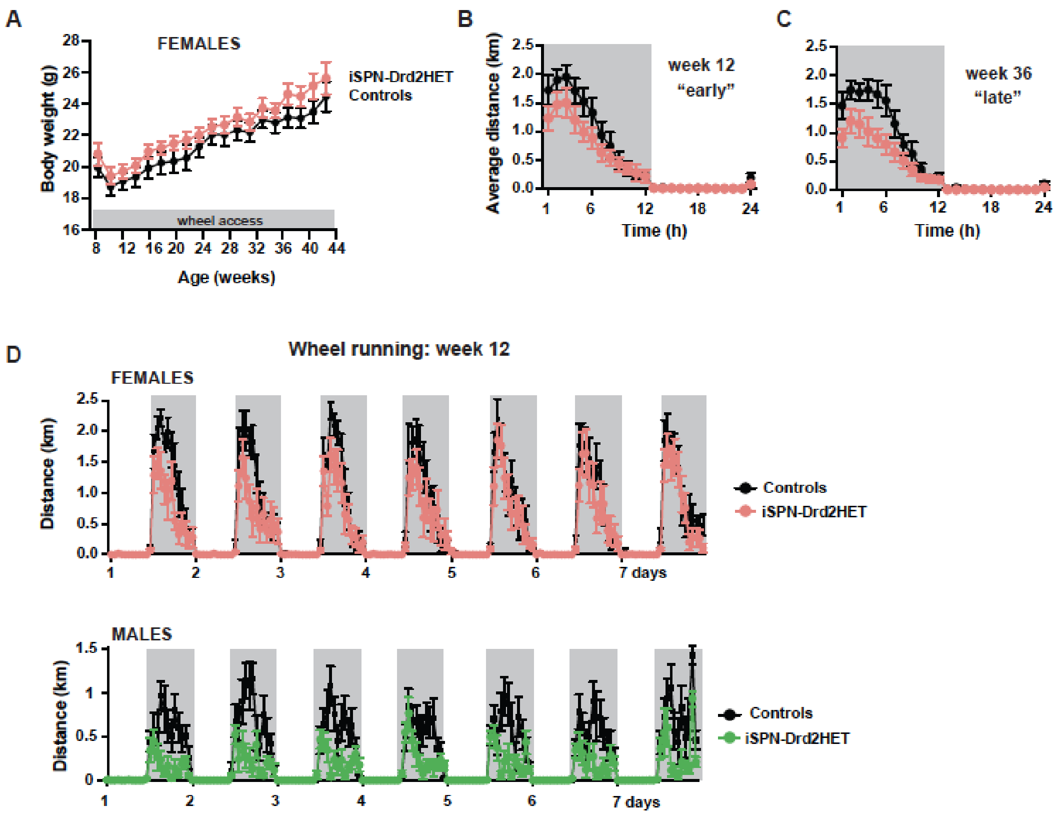


**Extended Data Figure 2. Female mice with a low density of D2 receptors in striatal neurons do not show significant differences in body weight and wheel running**

**A.** Longitudinal measure of body weight for female iSPN-Drd2HET (pink) and control (black) mice housed with a running wheel from week 8 - 44. Symbols and lines are mean ± SEM. **B-C**. Running distance per hour across the circadian cycle for iSPN-Drd2HET (pink) and control (black) female mice during (**B**) week 12 and (**C**) week 36. Symbols and lines are mean ± SEM. **D**. Running behavior over a week shows the circadian fluctuations in distance running for (top) female mice littermate $Drd2^{LoxP/wt}$ control (black) and iSPN-Drd2HET (pink) and for (bottom) male mice littermate $Drd2^{LoxP/wt}$ control (black) and iSPN-Drd2HET (green) mice. Symbols and lines are mean ± SEM.

**Extended Data Figure 3. Bocarsly et al.**

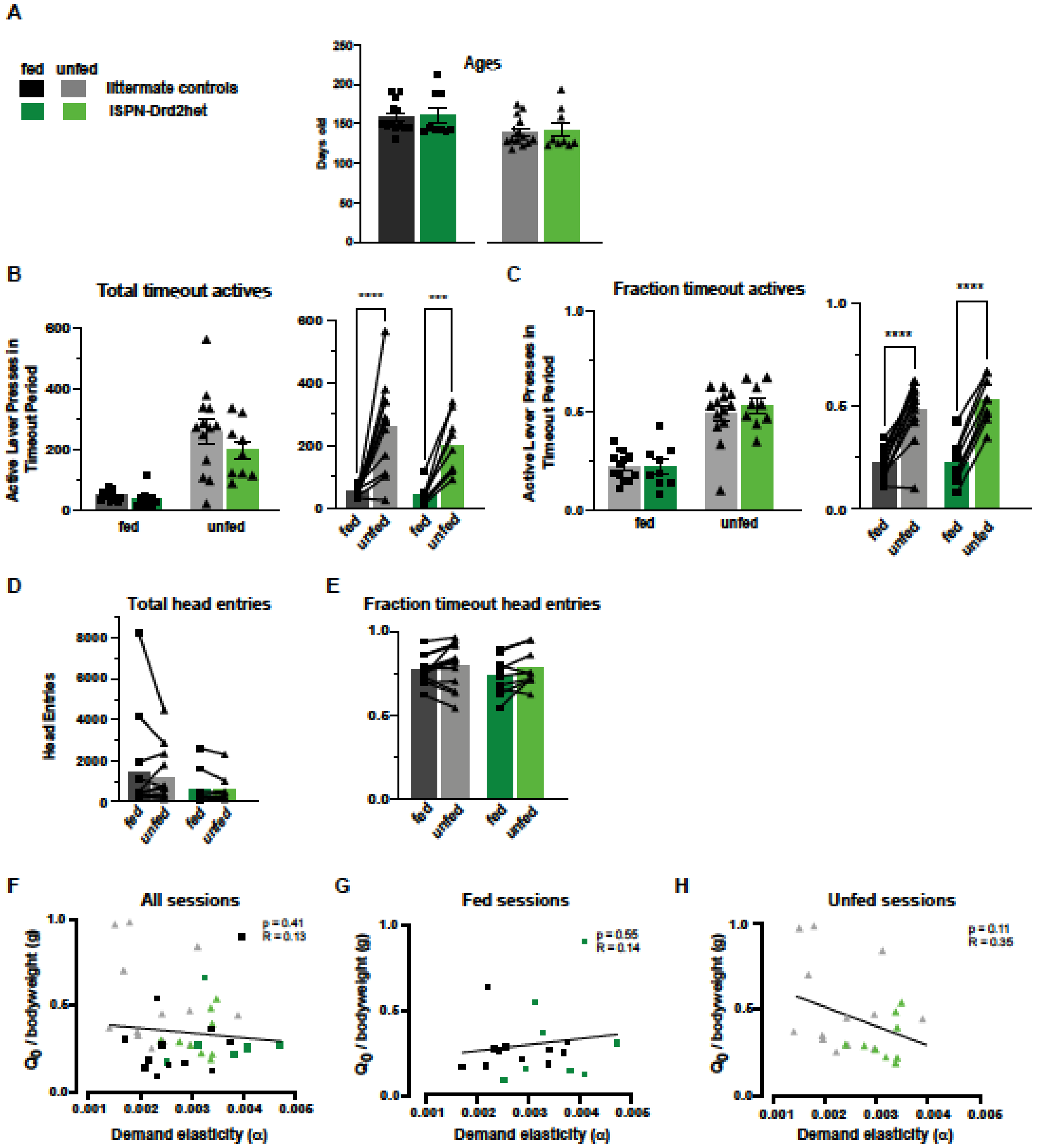


**Extended Data Figure 3. No difference in the age or persistent food-seeking behavior across genotypes, and independence of alpha and Q0 parameters**

**A.** Age (in days) of mice on the first day of behavioral economics testing in unfasted and fasted states. **B, C.** Total and fraction of active lever presses in timeout periods within behavioral economics sessions, showing summary (left) and paired (right) data. **D, E.** Total and fraction of timeout head entries within behavioral economics sections, showing paired data within mice. **F, G, H.** Alpha and Q0 fail to show a correlation, indicating the independence of these parameters. Symbols and lines represent mean ± SEM.

Extended Data Figure 4. Bocarsly et al.

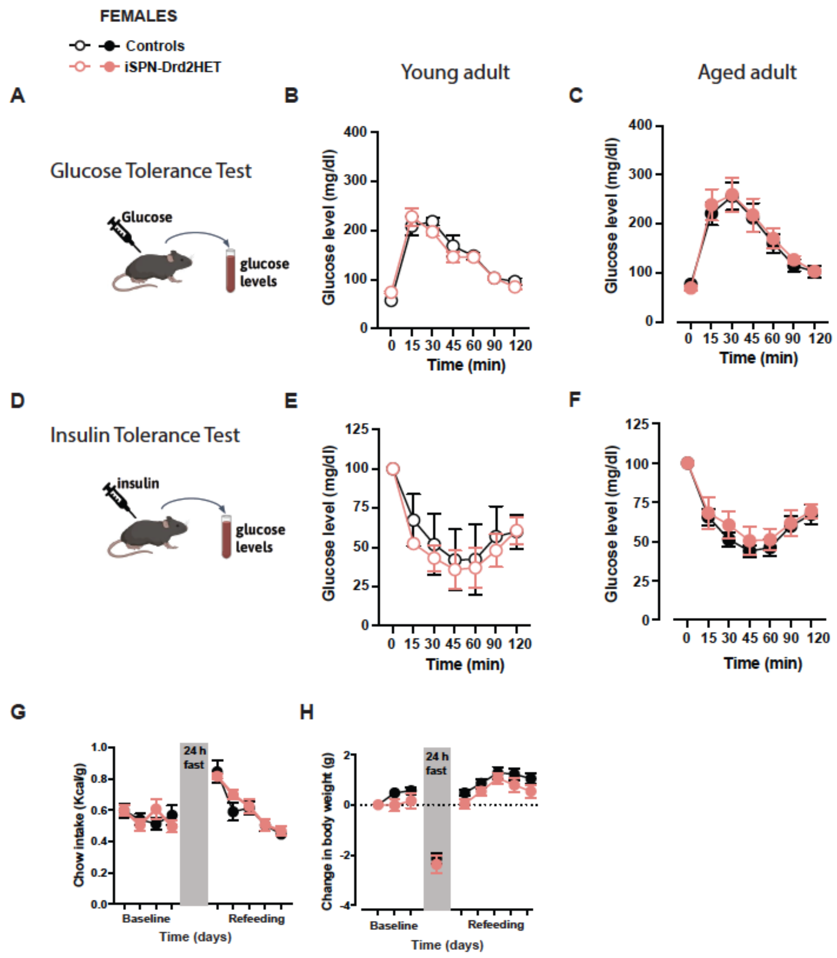


**Extended Data Figure 4. Metabolic signs of peripheral insulin resistance in aged overweight female mice**

All testing was performed in female young adult (8 weeks, left) and aged adult (60 weeks, right) iSPN-Drd2HET (pink) and control (black) mice. **A.** Schematic of i.p. glucose tolerance test. Mice were fasted overnight, administered an i.p. bolus of glucose, and blood glucose was monitored for 2 hours. **B.** Female mice were tested as young adults and **C.** aged adults. **D.** Similarly, on the i.p. insulin tolerance test, mice were fasted, however, this time they were administered an i.p. bolus of insulin, and blood glucose was monitored for 2 hours. As above, female mice were tested as **E.** young adults and **F.** aged adults. **G.** Chow intake and **H.** bodyweight were measured before and after a 24 h fasting period. No differences were seen between genotypes. For all panels, symbols and lines represent mean ± SEM.

Extended Data Figure 5. Bocarsly et al.

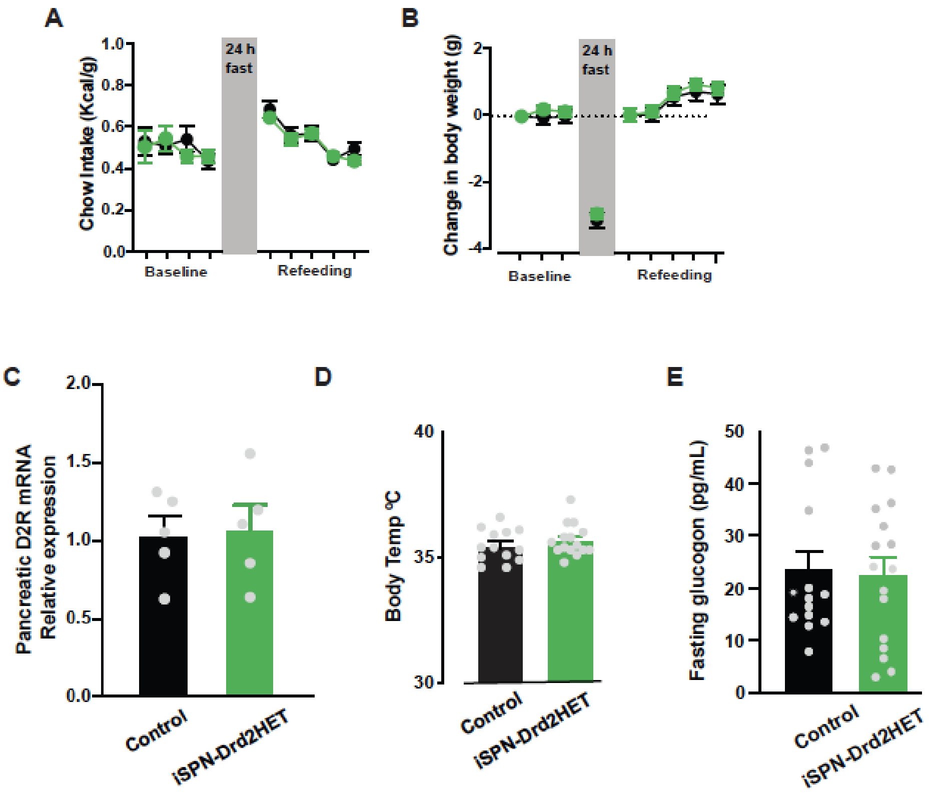


**Extended Data Figure 5. Not all peripheral markers of metabolism are changed in overweight male mice.**

**A.** Food intake and **B.** body weight were no different before and after a fasting re-feeding task in adult iSPN-Drd2HET mice and littermate controls. **C.** Drd2 mRNA levels were no different in pancreatic islets in male iSPN-Drd2HET mice and littermate controls. **D.** Basal body temperature was no different between male iSPN-Drd2HET mice and littermate controls. **E.** Fasting blood glucagon levels were no different between male iSPN-Drd2HET mice and littermate controls. All data is mean ± SEM.

**Extended Data Figure 6. Bocarsly et al.**

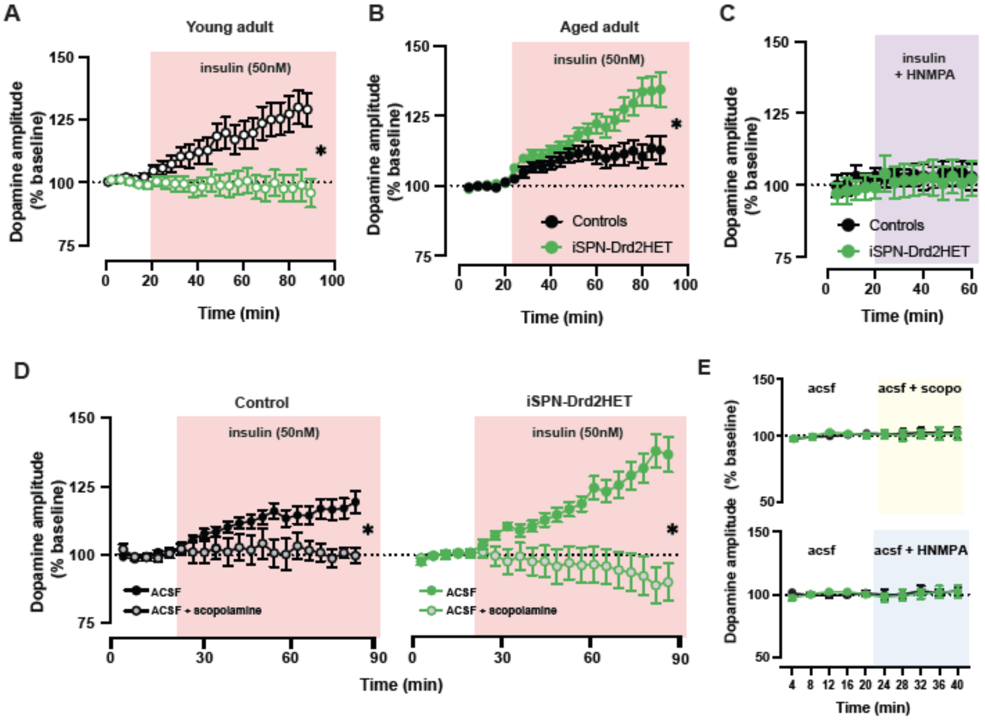


**Extended Data Figure 6. Time course of *ex vivo* dopamine response in young and aged male mice with low D2 receptors.**

**A.** In early adulthood (8 weeks) ex vivo FSCV was completed in fasted male iSPN-Drd2HET mice and littermate controls. When slices were treated with 50nM insulin, an increase in evoked dopamine was seen in control mice but absent in mice with low D2 receptors. **B.** When the same experiment was performed using tissue from 60-week-old, aged mice, while controls still showed an increase in evoked dopamine in response to insulin, mice with low D2 receptors showed an enhanced dopamine response to insulin. **C.** Control experiments demonstrate that the insulin receptor blocker, HNMPA, occludes the dopamine response to insulin in both iSPN-Drd2HET mice and littermate controls. **D.** Likewise, bath application of the acetylcholine muscarinic receptor blocker, scopolamine, blocks the dopamine response to insulin in both iSPN-Drd2HET (right) mice and littermate controls (left). **E.** Neither HNMPA nor scopolamine had any effects on evoked dopamine levels in the absence of insulin. Data is presented as mean ± SEM. * denotes statistical significance $p < 0.05$.
